# Membrane Elastic Properties During Neural Progenitor/Neural Stem Cell Differentiation

**DOI:** 10.1101/2020.04.03.019950

**Authors:** Juliana Soares, Glauber R. de S. Araujo, Cintia Santana, Diana Matias, Vivaldo Moura-Neto, Marcos Farina, Susana Frases, Nathan B. Viana, Luciana Romão, H. Moysés Nussenzveig, Bruno Pontes

## Abstract

Neural stem cells differentiate into several cell types that display distinct functions. However, little is known about how cell surface mechanics vary during the differentiation process. Here, by precisely measuring membrane tension and bending modulus, we map their variations and correlate them with changes in cell morphology along differentiation into neurons, astrocytes and oligodendrocytes. Both neurons and undifferentiated cells reveal a decrease in membrane tension over the first hours of differentiation followed by stabilization, with no change in bending modulus. Astrocytes membrane tension initially decreases and then increases after 72h, accompanied by consolidation of GFAP expression and striking actin reorganization, while bending modulus increases following observed alterations. For oligodendrocytes, the changes in membrane tension are less abrupt over the first hours but their values subsequently decrease, correlating with a shift from O4 to MBP expressions and a remarkable actin reorganization, while bending modulus remains constant. Oligodendrocytes at later differentiation stages show membrane vesicles with similar membrane tension but higher bending modulus as compared to the cell surface. Altogether, our results display an entire spectrum of how membrane elastic properties are varying, contributing to a better understanding of neural stem cell differentiation from a mechanobiological perspective.

## INTRODUCTION

The surface of all mammalian cells is composed of the plasma membrane cushioned underneath by a cortical actomyosin cytoskeleton. This pair of structures forms the membrane-cytoskeleton complex, a key regulator of several cellular processes, ranging from shape control and cell migration to molecule presentation and signalling [1,2]. The plasma membrane is the direct interface between the cytoplasm and the extracellular matrix [3,4], while the cortical cytoskeleton, also known as cell cortex, is a dynamic actomyosin meshwork that gives support to the plasma membrane [1,2]. The membrane-cytoskeleton complex exerts and reacts against forces owing to its elastic properties [1,2,4]. For brevity, we shall just refer to this complex as “cell membrane” (CM).

Over the years, different micromanipulation tools have been employed to exert forces on CMs so as to characterize their elastic responses [5]. Membrane tether pulling assays using optical tweezers (OT) [6,7] or atomic force microscopy (AFM) [8,9] have been used to extract nanotubes or tethers from CMs in order to determine these properties. By measuring both the equilibrium force and tether radius, the cell membrane surface tension (CMT - *σ*_*eff*_) and cell membrane bending modulus (CMBM - *K*_*eff*_) have been determined for different cell types [7,10-15]. CMT and CMBM result from joint contributions of cytoskeleton arquitecture, membrane composition and membrane-cytoskeleton attachment [14]. Moreover, these elastic properties (particularly CMT), as well as their changes, have been characterized as important regulators of cellular behaviors, specially regarding shape changes and force production [14,16]. Furthermore, it has been shown [13] that membrane elastic properties are correlated to cell function.

During development, neural stem cells, as neural progenitors (referred to just as NSCs) give rise to all neurons of the mammalian Central Nervous System (CNS). NSCs are also the source of two types of macroglial cells, the astrocytes and oligodendrocytes [17]. These three cell types are extremely different, not only in their morphological features but also in their functions. Neurons are highly anisotropic cells, with relatively quiescent and compact cell bodies (soma) containing the cell nucleus and dynamic protrusions (axons and dendrites), both susceptible to large structural changes [18]. Astrocytes are remarkably dynamic, constantly modifying their morphology during migration or when interacting with neurons [19]. Oligodendrocytes extend many protrusions that can ultimately form myelin sheaths, wrapping around axons to promote fast saltatory conduction of action potentials [20]. Several studies have revealed the molecular mechanisms that govern the path of NSCs differentiation and fate specification. Diverse growth factors and cytokines, together with epigenetic alterations including DNA methylation, histone modifications, and non-coding RNAs have already been described as key elements in this regard [21], as well as gene expression changes [22]. However, little is known about how the CM elastic properties of NSCs vary over the course of their differentiation nor the correlation between such variations and the final morphological phenotype of differentiated cells.

In the present work, we combined OT-based tether extraction experiments with measurements of tether radius using scanning electron microscopy (SEM), as well as fluorescence microscopy observations, to investigate the roles of the cytoskeleton on the CM elastic properties of NSCs along their differentiation into neurons, astrocytes and oligodendrocytes. Our aim was to map and collect experimental evidences that may provide a basis for future studies on how modifications in CM elastic properties correlate with shape and phenotype changes, ultimately influencing NSC differentiation.

## MATERIALS AND METHODS

### Animals

For NSC cultures, embryonic (E14; 14-day embryonic) swiss mice, obtained from pregnant females, were used. Mice were maintained at the Institute of Biomedical Sciences, Federal University of Rio de Janeiro. Animal husbandry and experimental procedures were performed in accordance with and approved by the Ethics Committee of the Health Sciences Center, Federal University of Rio de Janeiro (Protocol Number: 001-16).

### Cell Cultures

Primary cortical NSC cultures were prepared according to a modified protocol [23,24], using 14-days embryonic swiss mice. Briefly, NSCs suspensions were obtained by dissociating cells from the cerebral cortex and plated in Dulbecco’s Modified Eagle’s (DMEM) F-12 medium containing 0,6% glucose, N2, G5 (with FGF and EGF) and B27 supplements, 2mM L-glutamine, 5mM HEPES, 0,11% NaHCO_3_, and 1% penicillin/streptomycin (all from Invitrogen, Thermo Fisher, Carlsbad, CA, USA). The NSCs were cultured as neurospheres for 5 days. Then, they were mechanically dissociated and plated onto coverslips previously coated with 0.01% poly-L-lisine (Sigma-Aldrich, St. Louis, MO, USA). Five different cell conditions were used: (1) NSCs in neurospheres, (2) dissociated NSCs, (3) neurons, (4) astrocytes and (5) oligodendrocytes. All cells were obtained from NSCs that were induced to differentiate with specific media, partially replaced every 3 days, for 10 days (240h) under optimal culture conditions (37 °C and 5% CO_2_). All experiments were carried out at the following time points: 2, 24, 48, 72, 96, 120, 168 and 240h. NSCs in neurospheres and dissociated NSCs were maintained in their specific media. NSCs induced to differentiate into neurons were placed in Neurobasal media supplemented with 2mM L- glutamine, 1% penicillin/streptomycin and B27 supplement. NSCs induced to differentiate into astrocytes were placed in DMEM-F12 supplemented with 2mM L-glutamine, 10% fetal bovine serum and 1% penicillin/streptomycin. NSCs induced to differentiate into oligodendrocytes were placed in DMEM-F12 supplemented with 2mM L-glutamine, 0.5% fetal bovine serum, B27, 50μM T3, 5 μg/ml Insulin, 5 μg/ml transferrin, 5 ng/ml sodium selenite and 1% penicillin/streptomycin. All reagents, unless otherwise mentioned, were purchased from Invitrogen-Thermo Fisher Scientific (Carlsbad, CA, USA).

### Confocal Fluorescence Microscopy

Confocal fluorescence microscopy was performed for all the cell types and time points used in this study. Briefly, cells were fixed in PBS-paraformaldehyde 4% for 15 min, permeabilized with PBS-triton X100 0.2% for 5 min, blocked with PBS-BSA 5% (Sigma-Aldrich, St. Louis, MO, USA) for 1 hour and then incubated overnight at 4°C with primary antibodies for each of the specific markers: for NSCs, polyclonal antibody against brain lipid binding protein (BLBP) (Millipore, Merck KGaA, Germany), mouse antibody against nestin (Millipore, Merck KGaA, Germany), polyclonal antibody against sox2 (Invitrogen, Thermo Fisher, Carlsbad, CA, USA); for neurons, monoclonal antibody against β-tubulin III (Promega Corporation, Madison, WI, USA); for astrocytes, polyclonal antibody against glial fibrillary acidic protein (GFAP) (Dako, Denmark) and for oligodendrocytes, monoclonal antibody against oligodendrocyte marker O4 (R&D Systems, Minneapolis, MN, USA) and polyclonal antibody against myelin binding protein (MBP) (Abcam, UK). Then, secondary monoclonal and/or polyclonal Alexa antibodies conjugated with 546nm, 568nm or 633nm fluorophores (Molecular Probes Inc, Eugene, OR, USA) were incubated for 2h together with phalloidin-FITC (Molecular Probes Inc, Eugene, OR, USA). Coverslips were mounted on microscopy slides and visualized with a HC PL APO 63x/1.40 Oil CS objective lens attached to a Leica TCS-SP5 II confocal microscope (Leica Microsystems, Germany). Images were acquired using the LAS AF 2.2.0 Software (Leica Microsystems, Germany). Quantification analysis of F-actin and GFAP cytoskeleton networks was performed using FibrilTool [25], an ImageJ (National Institutes of Health, USA) plug-in capable of determining the average orientation of a fiber array, providing quantitative information about its anisotropy. The anisotropy value ranges from a maximum of 1, when all fibers point to the same direction in the array, to a minimum of 0, when they are all randomly oriented. Fluorescence quantifications for GFAP, MBP and O4 were performed using ImageJ. Briefly, an outline was drawn around each cell and the values for area, the mean grey (fluorescence) value and integrated density (*ID*) were obtained along with the corresponding measurements for the adjacent background. Then, the corrected total cell fluorescence (CTCF) was calculated for each experimental situation using the following equation:

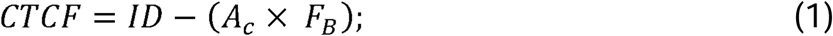

where *A*_*c*_ is the selected cell area and *F*_*B*_ is the mean grey (fluorescence) value of the background.

For GFAP, direct CTCF values were plotted for all experimental conditions. For O4 and MBP, the ratios between O4 CTCF and MBP CTCF were plotted for all experimental conditions.

### Optical Tweezers Setup and Calibration

The optical tweezers (OT) system employed an infrared Ytterbium linearly polarized colimated laser beam with a wavelength of 1064 nm and maximum power of 5W (model YLR-5-1064-LP) (IPG Photonics, NY, USA). The laser was coupled to an inverted Nikon Eclipse TE300 microscope (Nikon, Melville, NY, USA) equipped with a PLAN APO 100X 1.4 NA DIC H Nikon objective, used to create the trap. The OT system was calibrated using the same procedure previously described [13,26,27]. The trap transverse stiffness per unit power at the objective entrance was k/P = (0.25 ± 0.01) pNμm^−1^mW^−1^.

### Tether Extraction Experiments with Optical Tweezers

Tether extraction experiments using optical tweezers were performed following the same procedures previously described [7,10-13]. Briefly, NSCs in neurospheres and/or dissociated NSCs were plated and allowed to attach to glass bottom dishes. NSCs in neurospheres were allowed to attach only for 2h prior to experiments. Dissociated NSCs were allowed to attach and differentiate for all time points, as described above. Then, uncoated polystyrene beads (radius = 1.52 ± 0.02 µm) (Polysciences, Warrington, PA) were added and each one of the glass bottom dishes containing the cells was placed in the OT microscope. A bead was trapped and pressed against a chosen cell for ∼5 seconds, allowing its attachment to the cell surface. The microscope motorized stage (Prior Scientific, Rockland, MA, USA) was then set to move with controlled velocity (1 µm/s). Movies were collected at a frame rate of 10 frames/second using a Hamamatsu C2400 CCD camera (Hamamatsu, Japan) coupled to a SCION FG7 frame grabber (Scion Corporation, Torrance, CA, USA). Using the trap calibration and the measured bead position displacement, obtained by analysis of images extracted from the movie, the tether force, *F*_0_, was determined. All experiments in the OT microscope were conducted in optimal culture conditions (37°C and 5% CO_2_). Data analysis and force calculations were performed using ImageJ and Kaleidagraph (Synergy Software, Essex Junction, VT, USA) softwares.

### Measurements of Tether Radii

After extracting the cell tethers, the beads used during the extraction were attached to the coverslip. The samples were then fixed and prepared for SEM following the same procedures previously described [7,10-13]. Thus, the mean values for tether radii, *R*, corresponding to each experimental situation were obtained.

However, the aforementioned procedure could not be employed to measure the tether radii extracted from oligodendrocyte vesicles. To circumvent this issue we adopted another previously validated method, already tested for the same purpose [7,13]. This method is based on the force barrier theory for tether formation [28] and its theory gives the following relation:

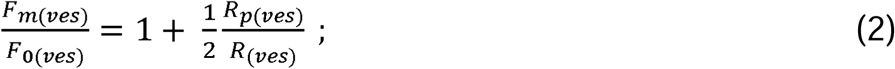

where *F*_0(*ves*)_ is the vesicle tether force, *F*_*m*(*ves*)_ is the vesicle maximum force before tether formation, *R*_(*ves*)_ is the vesicle tether radius and *R*_*p*(*ves*)_ is the radius of the circular patch contact area between the bead and the vesicle membrane.

The values of *F*_*m*(*ves*)_ and *F*_0(*ves*)_ were experimentally determined; *R*_*p*(*ves*)_ was measured from the image of the bead and the deformed vesicle surface. The value of *R*_(*ves*)_ was then obtained from equation (2).

### Determination of Membrane Elastic Properties

Once the values of *F*_0_ and *R* were measured, it was then possible to determine the values of the cell membrane tension (CMT - *σ*_*eff*_) and the cell membrane bending modulus (CMBM - *K*_*eff*_) [29,30].

The CMT (*σ*_*eff*_) is given by:

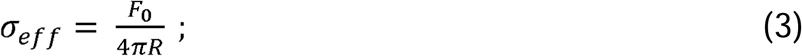

and the CMBM (*K*_*eff*_):

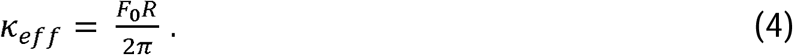

The values of membrane tension, *MT*_(*ves*)_, and bending modulus, *BM*_(*ves*)_, for oligodendrocyte vesicles’ were also determined using the equations above.

### Statistical Analysis

Some data are presented as mean ± standard error. For tether force and radius values, Box-and-Whiskers plots were used. The boxes extend from the 25th to 75th percentiles, with a black horizontal line at the median and a black cross at the mean; black whiskers extend from 5th to 95th percentiles for all experimental conditions; values outside these ranges are plotted as individual points. Data were analyzed using GraphPad Prism statistics software (GraphPadSoftware, Inc. La Jolla, CA, USA). Mann-Whitney *U*-tests were used for comparisons between each situation and the 2h condition. *means *p* < 0.05; ** means *p* < 0.01; *** means *p* < 0.001 and **** means *p* < 0.0001. The p-values and other numbers for all experiments are provided in the figure legends.

## RESULTS

### OT as a screening tool to measure membrane elastic properties of differentiated and undifferentiated NSCs

Aiming to characterize the CM elastic properties of NSCs induced to differentiate into astrocytes, oligodendrocytes and neurons and to compare the results with those found for cells maintained with their stem capacity, either as neurospheres or isolated cells, OT-based tether extraction experiments were performed for each cell type over periods of 2 to 240h in culture. As an example of the experimental procedures, images associated with a tether extraction experiment performed in a neuron differentiated cell are shown in Figure 1A (extracted tether is indicated by a white arrow) and Figure 1B (zoom of the extracted tether). The corresponding tether extraction force curve is shown in Figure 1C. The tether force, *F*_0_, (referred to as the steady-state force in Figure 1C) and the tether radius, *R*, (Figures 1D – F) were carefully measured for each experimental time point. The results are presented in the following sections.

**Figure 1:**
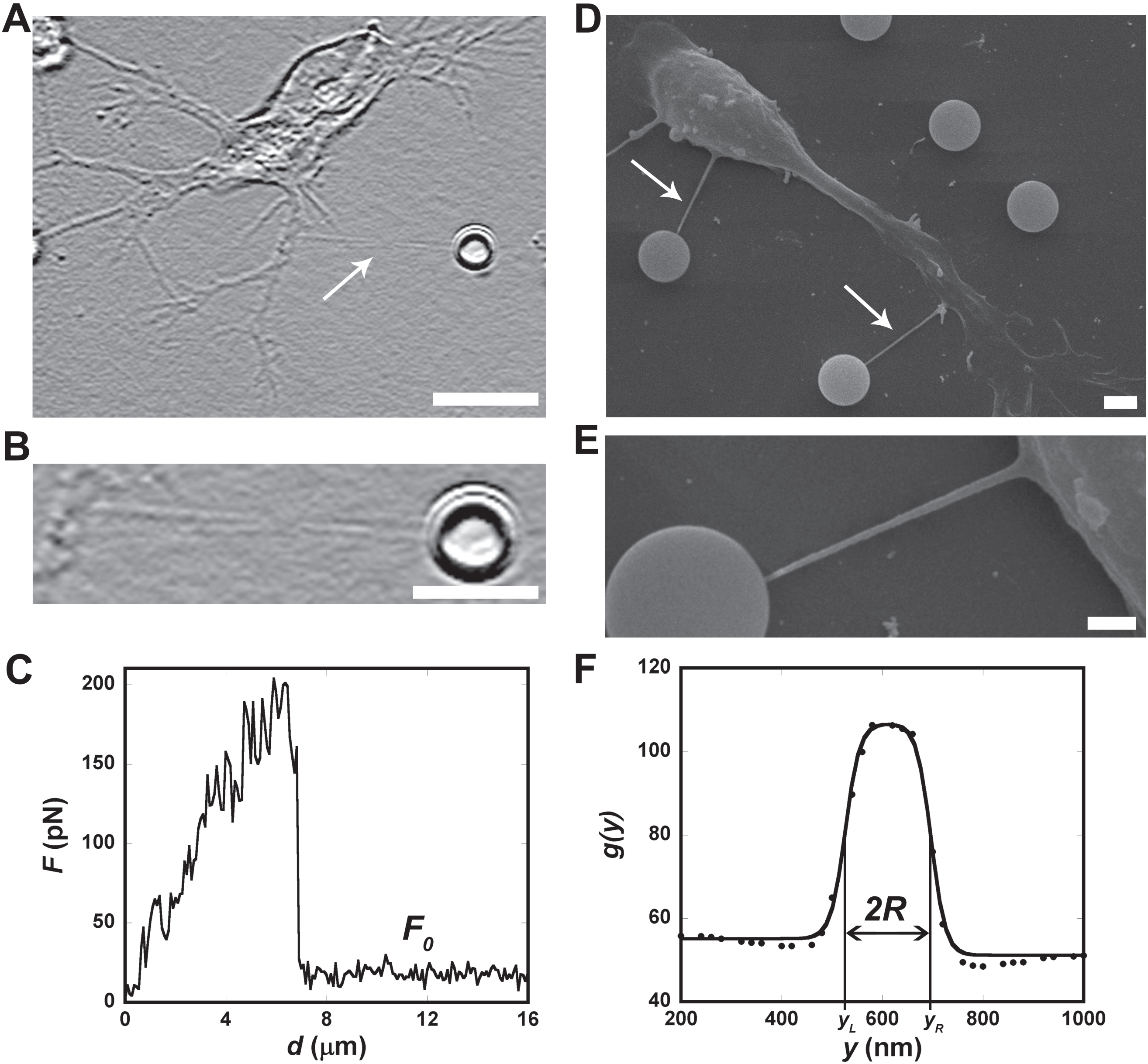
Typical tether force and radius measurements. (A) Representative bright field image of a 48h-differentiated neuron extracted tether (indicated by a white arrow). ImageJ shadow north processing filter was applied to better visualize the tether. (B) Zoom of the tether in A. Scale bar for A is 10 µm and for B is 5 µm. (C) A typical tether extraction force curve. *F*_0_ is the steady-state tether force. (D) SEM representative image of tethers extracted from a typical 24h-differentiated neuron. (E) Zoom of one of the tethers indicated by the white arrows in D. Scale bar for D is 2 µm and for E is 1 µm. (F) Grey level plot profile of the tether in E.

### Membrane elastic properties measured for undifferentiated neural stem cells vary only in the initial hours after plating

NSCs, previously grown as neurospheres for 5 days (Figure S1A), were dissociated and replated as isolated cells in stem cell medium to maintain their stemness (Figure S1B). Their morphologies and the expression of specific markers, were followed from 2 to 240h after plating. Figure 2A displays the actin cytoskeleton of these cells (stained in green for phaloidin-FITC, a molecule commonly used as a cytochemical marker of polymerized actin (F-actin)) together with a specific marker, the brain lipid binding protein (BLBP or FABP7 - stained in red), known to be expressed in radial glial cells during development. It has been documented that radial glial cells not only participate as scaffolds that allow neurons to migrate within the developing cortex [31,32], but they also function as neural progenitors cells [33-35]. Thus, the NSCs used in this study (Figure 2A) have the morphology and characteristics of radial glial cells. Moreover, to double-check that these cells were kept undifferentiated throughout the experimental procedures, NSCs were also stained, after 240h in culture, for nestin (in white) and sox2 (in red) (typical stem cell markers) together with phaloidin-FITC (in green) and the cell nucleus, stained with dapi (in blue) (Figure S1B).

**Figure 2:**
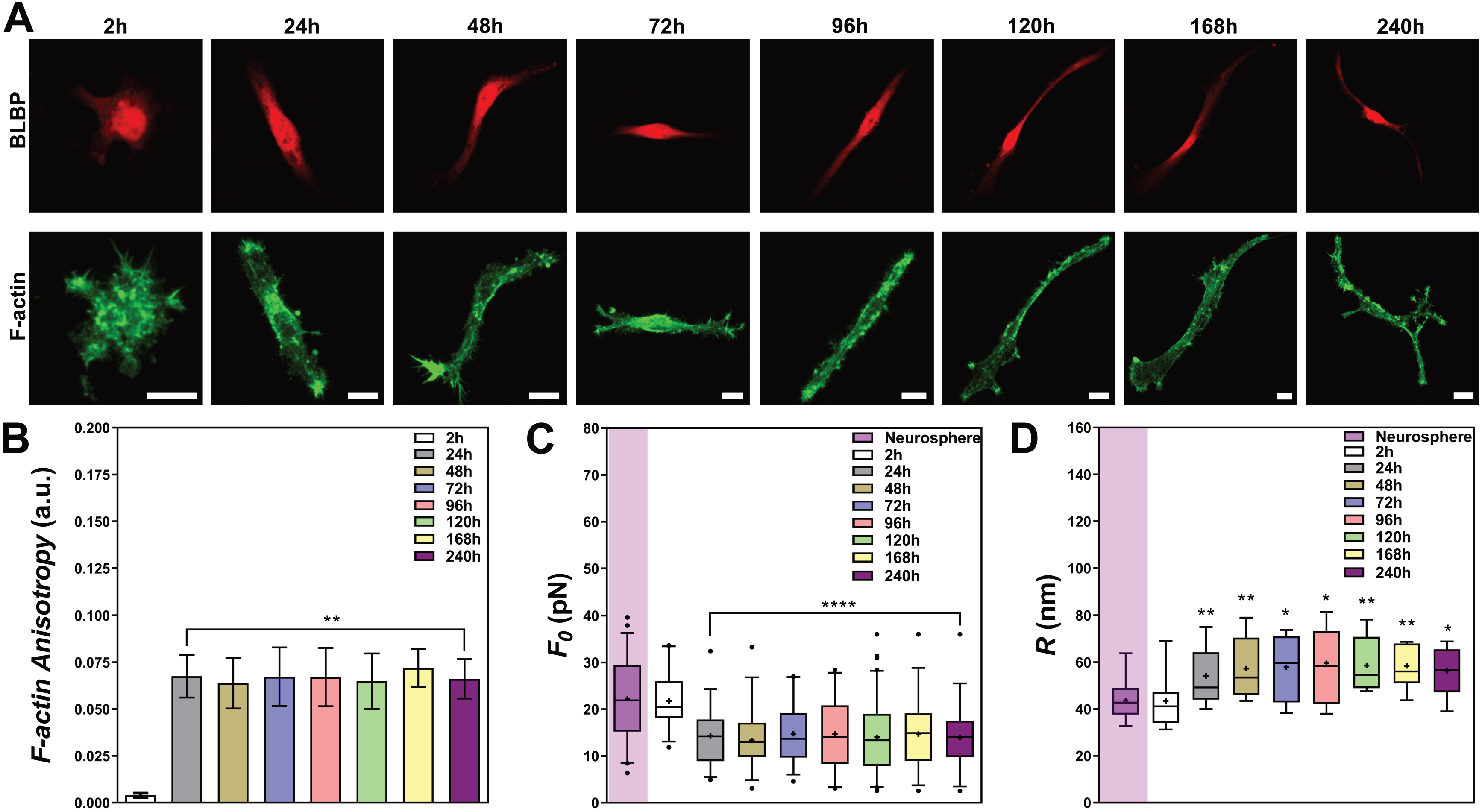
Tether extraction experiments and radii measurements for undifferentiated neural stem cells. (A) Representative images of undifferentiated NSCs stained for BLBP (red) and F-actin, with phaloidin-FITC, (green) from 2h to 240h. Scale bars are all 10 μm. (B) Plot of the mean anisotropy values of F-actin after FibrilTool analysis. At least 10 different cells for each differentiation time condition were imaged and analyzed. They all show similar behaviors as the ones represented in A. Standard errors were used as error bars. (C-D) Plot of tether force values (at least 32 different measurements) (C) and tether radii values (at least 12 different measurements) (D) for each experimental group. The colored boxes extend from 25th to 75th percentiles, with a black horizontal line at the median and a black cross at the mean; black whiskers extend from 5th to 95th percentiles; values outside these ranges are plotted as individual points. Tether force and radius values for cells in neurospheres are also represented (highlighted with a purple rectangle) for comparison purposes. * means *p* < 0.05, ** means *p* < 0.01, **** means *p* < 0.0001 in *Mann-Whitney U-*test statistics when comparing each time point with the 2h condition.

The actin cytoskeleton architecture, with the exception of the 2h condition, appeared to not present any substantial variations throughout the days of culture, as verified by the images (Figure 2A). However, to better quantify these visual observations, we also performed a quantitative analysis using FibrilTool [25]. The results of this analysis (Figure 2B) showed that, overall, the cells present a similar F-actin anisotropy throughout the entire experiment, except for the cells at the 2h time point.

Next, tether extractions were performed on dissociated NSCs that were maintained undifferentiated in culture. The corresponding mean values of tether force are shown in Figure 2C. No statistically significant change in tether force values were found for NSCs from 24 to 240h in culture (Figure 2C). However, a ∼1.5-fold decrease in tether force value (from 22±1 to estimated 14±1 pN) was found when comparing the 2h condition with all the other ones (Figure 2C).

The tether radii were also measured using SEM. Figure 2D shows the mean values for the NSCs from 2 to 240h in culture. No statistically significant change in tether radii values were found for NSCs from 24 to 240h in culture (Figure 2D). However, a ∼1.3-fold increase in tether radii value (from 43±3 to estimated 57±4 nm) was found when comparing the 2h condition with all the other ones (Figure 2D).

In order to compare the CM elastic properties of dissociated NSCs versus NSCs in spheres, we cultured neurospheres for 5 days (Figure S1A – phase contrast images), plated and allowed them to attach for 2h in stem cell medium (Figure S1A - nestin (in white), sox2 (in red) and dapi (in blue)). Tether force and radii measurements were also performed on different cells around the neurospheres. The mean values of *F*_0_ and *R* are shown in Figures 2C and 2D, respectively. No statistically significant changes in tether force and radius values were found between the 2h-dissociated NSCs and those grown as neurospheres. However, the similar ∼1.5-fold decrease in tether force value and the ∼1.3-fold increase in tether radius were still evident when comparing those neurospheres condition with the 24-240h conditions.

Taken together, the results confirm that NSCs remain undifferentiated over the entire experiment, with an initial 2-fold drop in their CMT values in the very early hours, followed by stabilization in the subsequent hours (Figure 7A, grey curve and dots). The CMBM values, on the other hand, remain constant over time (Figure 7A, red curve and dots). These variations in CMT are not attributed to the dissociation of neurospheres but may be related to cell spreading and acquisition of final morphological phenotype.

### Membrane elastic properties measured for neurons show a similar pattern in comparison to undifferentiated neural stem cells

When dissociated, some NSCs were kept undifferentiated, but others were plated in a culture dish containing neurobasal medium, supplemented with factors that induce cells to differentiate into neurons. Again, the cell morphology and the expression of specific markers were followed from 2h to 240h after plating. Figure 3A shows the phaloidin-FITC labeled actin cytoskeleton (in green), together with β-tubulin III (in red), known to be expressed in neurons [36]. The results show that the cells acquire the morphology and characteristics of neurons, with several neurites and growth cones, which began to appear within the first 24h and increased in the subsequent hours of culture (Figure 3A), confirming that NSCs already differentiate into neurons after being in culture for only a few hours.

**Figure 3:**
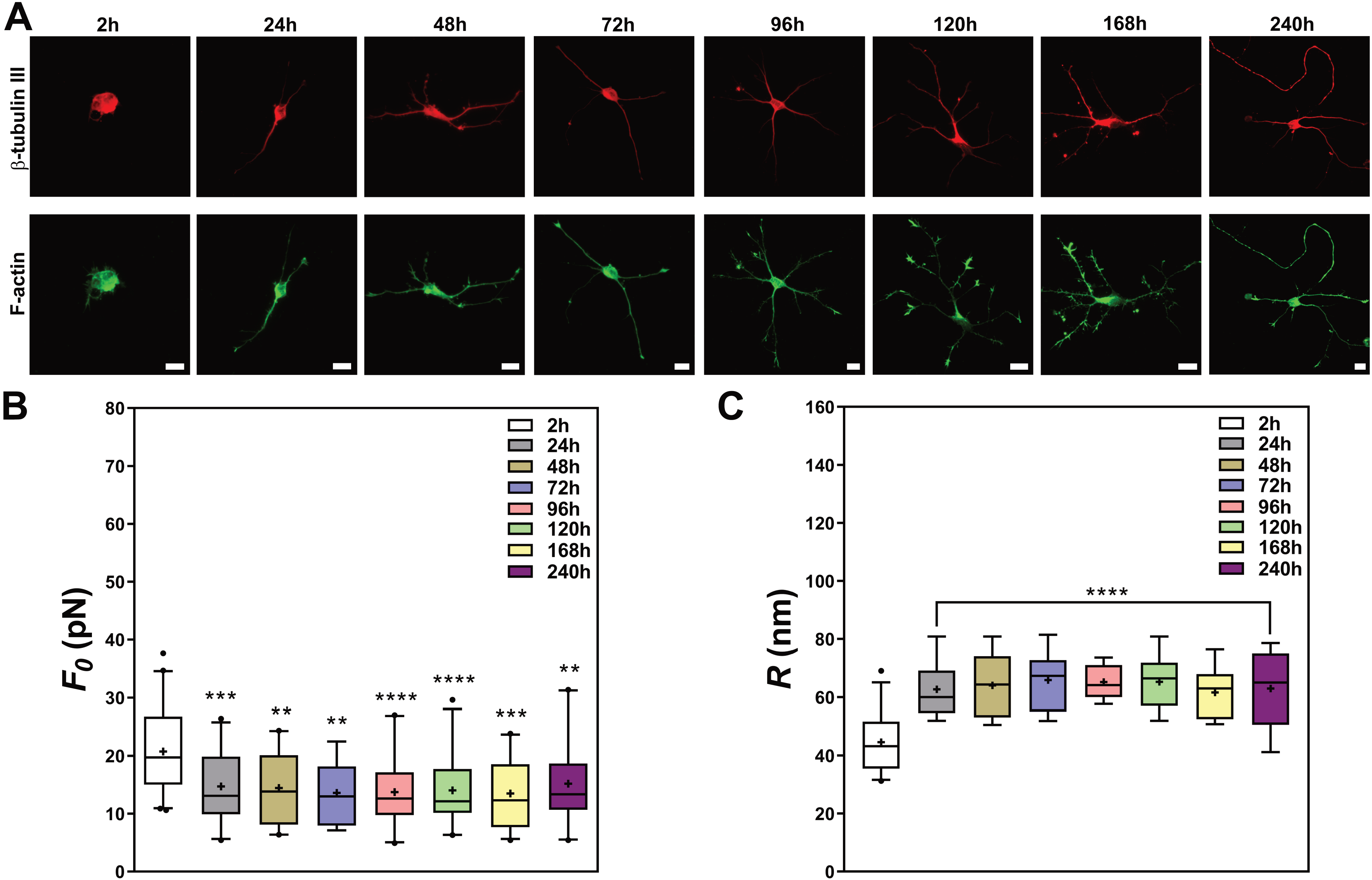
Tether extraction experiments and radii measurements for NSCs induced to differentiate into neurons. (A) Representative images of cells stained for β-tubulin III (red) and F-actin (green) from 2h to 240h. Scale bars are all 10 μm. (B-C) Plot of tether force values (at least 18 different measurements) (B) and tether radius values (at least 10 different measurements) (C) for each experimental time point. The colored boxes extend from 25th to 75th percentiles, with a black horizontal line at the median and a black cross at the mean; black whiskers extend from 5th to 95th percentiles; values outside these ranges are plotted as individual points. * means *p* < 0.05, ** means *p* < 0.01, *** means *p* < 0.001, **** means *p* < 0.0001 in *Mann-Whitney U*-test statistics when comparing each time point with the 2h condition.

OT-based tether extractions were performed on these neurons along the differentiation period. The mean values of *F*_0_ are shown in Figure 3B. No changes in tether force values were found for neurons from 24h to 240h in culture (Figure 3B). However, a ∼1.5-fold decrease in tether force values (from 21±1 to estimated 14±1 pN) was found when comparing the 2h condition with all other ones (Figure 3B).

Regarding the tether radii results, no statistically significant change was detected among the time points ranging from 24h to 240h in culture (Figure 3C). However, a ∼1.4-fold increase in tether radii values (from 45±2 to estimated 64±3 nm) was found when comparing the 2h condition with all other ones (Figure 3C).

In view of the prior demonstration that, regardless of the region (cell body, neurites and growth cones) there is no significant difference in the tether force and radius [13], we chose to pool all the measurement results together in the plots (Figures 3B and 3C). The values found for the tether force and radius were also of the same order of magnitude as those previously found for mouse cortical and ganglionic eminence neurons [13].

The results confirm that NSCs are able to differentiate into neurons with an initial 2.1-fold drop in their CMT values in the very early hours, followed by stabilization in subsequent hours, while increasing the number and length of neurites and number of growth cones over time. The CMBM values, on the other hand, do not vary during the experimental time points. Finally, the values for the CMT (Figure 7B, grey curve and dots) and CMBM (Figure 7B, red curve and dots) are very similar to those found for undifferentiated NSCs (Figure 7A). Again, variations in CMT may be related to cell spreading and acquisition of final morphological phenotype.

### Astrocyte differentiation process reveals interesting patterns that correlate cytoskeletal architecture remodeling with changes in membrane elastic properties

Dissociated NSCs were also induced to differentiate into astrocytes. Again, the cell morphology and the expression of specific markers were followed from 2h to 240h after plating. Figure 4A shows the actin cytoeskeleton, stained for phaloidin-FITC (in green), together with a specific marker for astrocytes, GFAP (in red) [37]. The results demonstrate that even though the cells acquired the morphology and characteristics of astrocytes, the expression of GFAP began showing up as filaments, in some cells, only 48h after plating, yielding more stable filaments after 72h of culture (Figures 4A and 4B), confirming that the NSCs differentiated into astrocytes. Interestingly, the actin cytoskeleton also presented a morphological shift, from small actin filaments (in the first 48h to 72h) to very well-defined stress fibers after 96h (Figure 4A). These results were confirmed by the FibrilTool analysis, which showed that the anisotropy of both actin and GFAP cytoskeletal networks increased with time in culture (Figures 4B and 4C).

**Figure 4:**
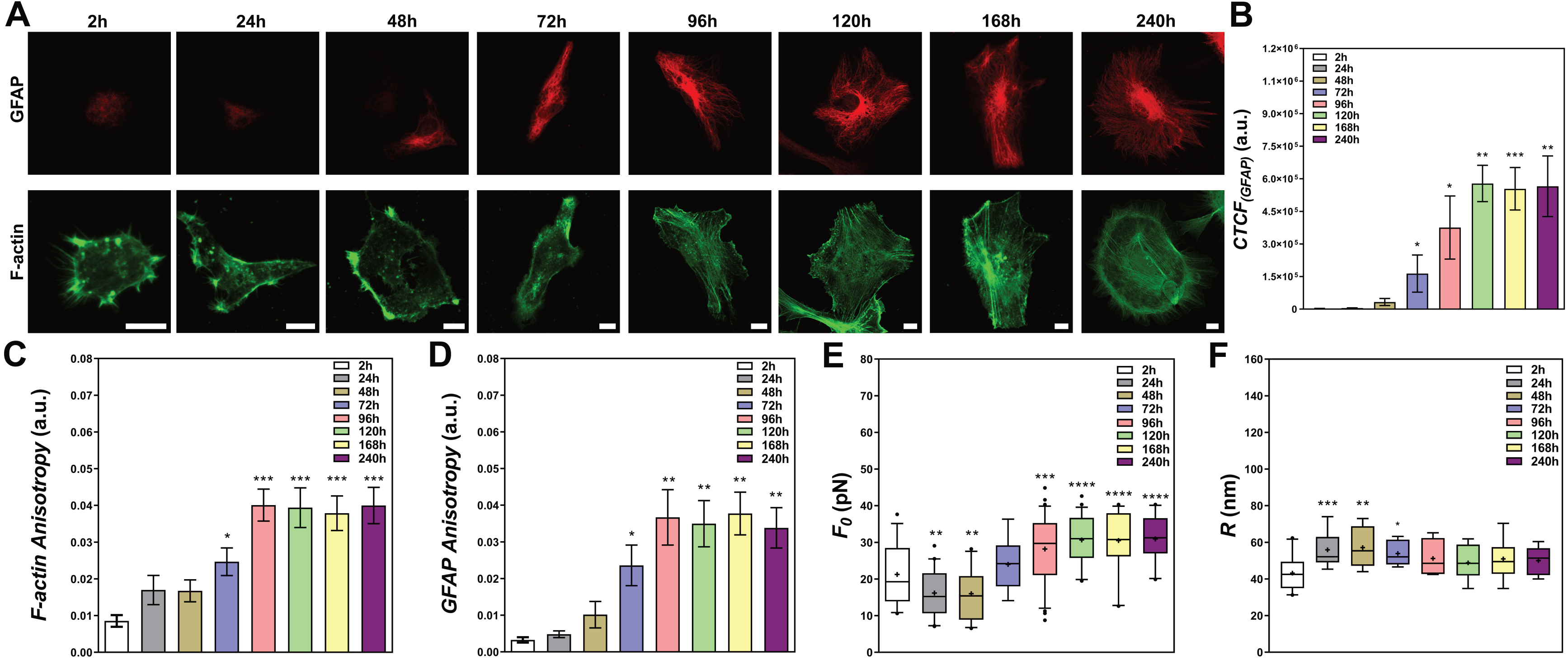
Tether extraction experiments and radii measurements for NSCs induced to differentiate into astrocytes. (A) Representative images of cells stained for GFAP (red) and F-actin (green) from 2h to 240h. Scale bars are all 10 μm. (B) Plot of the mean CTCF values for GFAP over the course of differentiation from 2h to 240h. (C-D) Plot of the mean anisotropy values of F-actin staining (C) and GFAP staining (D) after FibrilTool analysis. At least 10 different cells for each condition were analyzed. Standard errors were used as error bars in B, C and D. They all show similar behaviors as the ones represented in A. (E-F) Plots of tether forces (at least 16 different measurements) (E) and tether radii values (at least 10 different measurements) (F) for each time point. The colored boxes extend from 25th to 75th percentiles, with a black horizontal line at the median and a black cross at the mean; black whiskers extend from 5th to 95th percentiles; values outside these ranges are plotted as individual points. * means *p* < 0.05, ** means *p* < 0.01, *** means *p* < 0.001, **** means *p* < 0.0001 in *Mann-Whitney U*-test statistics when comparing each time point with the 2h condition.

Tether extraction measurements were performed in these astrocytes throughout their differentiation in culture. The mean values of tether force are shown in Figure 4D. A clear decrease (∼1.3-fold) in tether force was observed from 2h to 48h, followed by a slight recovery at 72h after plating, consistently correlated with the increase in GFAP expression and the shift in F-actin architecture described above (Figures 4A, 4B 4C and 4D). Moreover, the tether force values found after 96h (∼30 pN) increased∼1.4-fold when compared with the 2h result and also increased almost 2.0-fold when compared with the 24 – 48h conditions (Figure 4D).

Figure 4E shows the mean radii values measured for cells induced to differentiate into astrocytes. A clear increase (∼1,3-fold) in radii values from 24h to 72h and a slight increase from 96h to 240h (∼1.16-fold) when compared to 2h time point were also observed (Figure 4E).

Altogether, the results confirm that even though NSCs differentiate into astrocytes and remain differentiated, their differentiation process is different from the neuronal differentiation, since GFAP expression starts to appear only 48h after induction, whereas in neurons the expression of β-tubulin III occurs in the very first hours. The shift in actin cytoskeleton architecture together with the increase in GFAP expression are both correlated with the changes in CM elastic properties observed for astrocytes after 48 – 72h. The results show that there is a ∼1.3-fold increase in CMT (Figure 7C, grey curve and dots) and a 1.7-fold increase in CMBM (Figure 7C, red curve and dots) in differentiated astrocytes (astrocytes from 96 – 240h) when compared to the undifferentiated cells (Figure 7A).

The values of CMT and CMBM found for astrocytes at later differentiation stages are within the same order of magnitude of those previously found for differentiated astrocytes obtained from newborn mice [13].

### Oligodendrocyte differentiation reveals interesting patterns that correlate cytoskeletal remodeling and expression of specific markers with changes in membrane elastic properties

Finally, NSCs were also induced to differentiate into oligodendrocytes. Likewise, we followed the cell morphology and expression of specific markers during the differentiation process. Figure 5A shows the actin cytoeskeleton (in green), together with two specific markers for oligodendrocytes: O4 (in red) and MBP (in white) [38]. The results show that the cells acquired the morphology and characteristics of oligodendrocytes. Interestingly, the expression of O4 increased in the first hours of differentiation and then decreased after the 96h time point. Conversely, the expression of MBP, which was low in the first hours, increased in the last hours, especially after the 96h time point (Figure 5A). The antagonic expression patterns between O4 and MBP become more evident in Figure 5B, where the ratios between O4 and MBP fluorescence intensities were plotted. The cell morphology and actin cytoskeleton organization also dynamically changed over time in culture. Cells that were induced to differentiate into oligodendrocytes started to present a star-like branched morphology during the first 48h, with several F-actin containing protrusions (Figure 5A). However, this ramified morphology changed to a more flat-like lamellar shape, with the formation of membrane extensions, after 72 – 96h. These changes were correlated with the increase in MBP expression and the simultaneous decrease in O4 expression. Interestingly, in the same time frame, the actin cytoskeleton also presented peculiar changes in its architecture. These changes first appeared in the outermost region of the membrane extensions surrounding the cell as a sort of ring and then disappeared in the subsequent hours (168h and 240h) (Figure 5A). This morphological behavior had already been documented in other studies [39-41].

**Figure 5:**
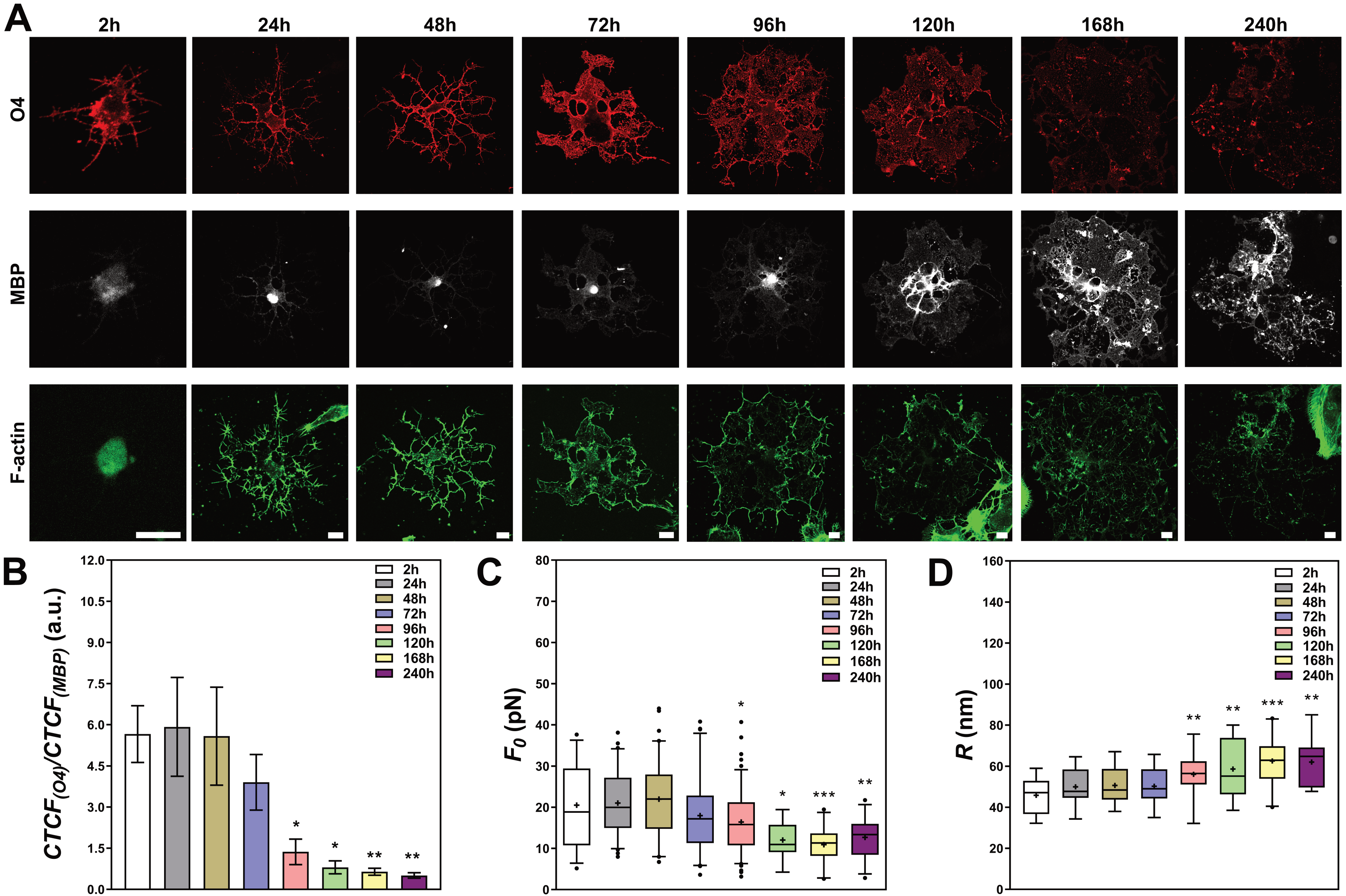
Tether extraction experiments and radii measurements for NSCs induced to differentiate into oligodendrocytes. (A) Representative images of cells stained for O4 (red), F-actin (green) and MBP (white) from 2h to 240h. Scale bars are all 10 μm. (B) Ratio of O4 and MBP CTCF values over the course of differentiation from 2h to 240h. At least 10 different cells for each condition were imaged and analyzed in B. They all show similar behaviors as the ones represented in A. (C-D) Plots of tether forces (at least 20 different measurements) (C) and tether radii values (at least 12 different measurements) (D) for each condition. The colored boxes extend from 25th to 75th percentiles, with a black horizontal line at the median and a black cross at the mean; black whiskers extend from 5th to 95th percentiles; values outside these ranges are plotted as individual points. * means *p* < 0.05, ** means *p* < 0.01, *** means *p* < 0.001 in *Mann-Whitney U*-test statistics when comparing each time point with the 2h condition.

Tether extraction experiments were performed in oligodendrocytes along their differentiation process. The mean values of *F*_0_ are shown in Figure 5C. No differences in tether force were observed in the first 72h as compared to the 2h time point (Figure 5C). However, a statistically significant decrease in tether force was observed in the time points between 96h and 240h (Figure 5C). This result can be well correlated with the increase in MBP expression, the change in oligodendrocyte morphology and the shift in actin cytoskeleton architecture described above (Figures 5A). The values for 120, 168 and 240h (on the order of ∼12pN of force) were ∼1.7 times lower that those found within the first 48h (on the order of ∼21pN of force).

Concerning the mean radii values, Figure 5D depicts how these numbers changed along the oligodendrocyte differentiation process. We observed a clear increase (∼1,3-fold) in tether radii from 120 to 240h when compared to the 2h condition (Figure 5D).

Another feature that appeared in the last hours of differentiation (168 – 240h) was the presence of spherical vesicle-like membrane protrusions steming from the oligodendrocytes surfaces (Figure 6A, images 1 and 2). These vesicles had been previously documented several years ago in ultrastructural caracterizations of cultured oligodendrocytes [42], but apparently neglected since that time.

**Figure 6:**
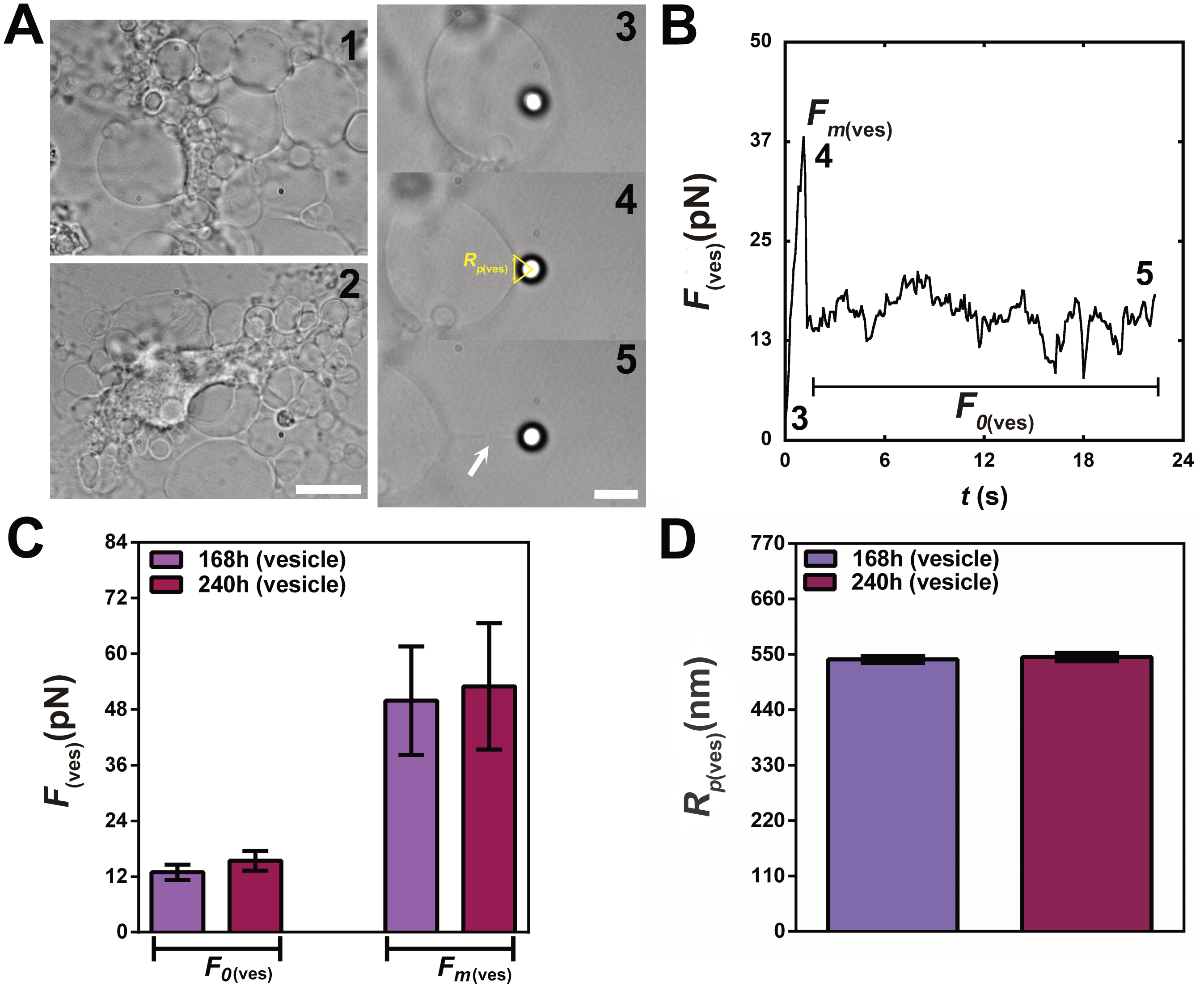
Tether extraction experiments and patch radii measurements for oligodendrocyte vesicles. (A) Typical bright field images of oligodendrocytes at 168h (Image 1) and 240h (Image 2) of differentiation, presenting several vesicles along their surfaces. (Images 3-5) Selection of images of a typical tether extraction experiment from the surface of oligodendrocyte vesicles. (Image 3) Initial moment, bead is being pressed against the oligodendrocyte vesicle with the optical trap, (Image 4) moment when the force reaches the maximum value – schematics indicating how *R*_*p*(*ves*)_ is obtained – and, (Image 5) membrane tether already formed, indicated as a white arrow. Scale bar for Images 1-2 is 10 μm and for Images 3-5 is 5 μm. (B) Force curve of a tether extracted from the oligodendrocyte vesicle. Numbers 3, 4, and 5 in the plot correspond to the images of the same numbers in A. *F*_*m*(*ves*)_ means the maximum force before tether formation and *F*_0(*ves*)_ means the steady-state tether force. (C) Plot of the mean values of *R*_*p*(*ves*)_ for each experimental condition, as indicated by the plot legends. Standard errors were used as error bars in C and D. At least 20 different experiments were performed for each situation.

**Figure 7:**
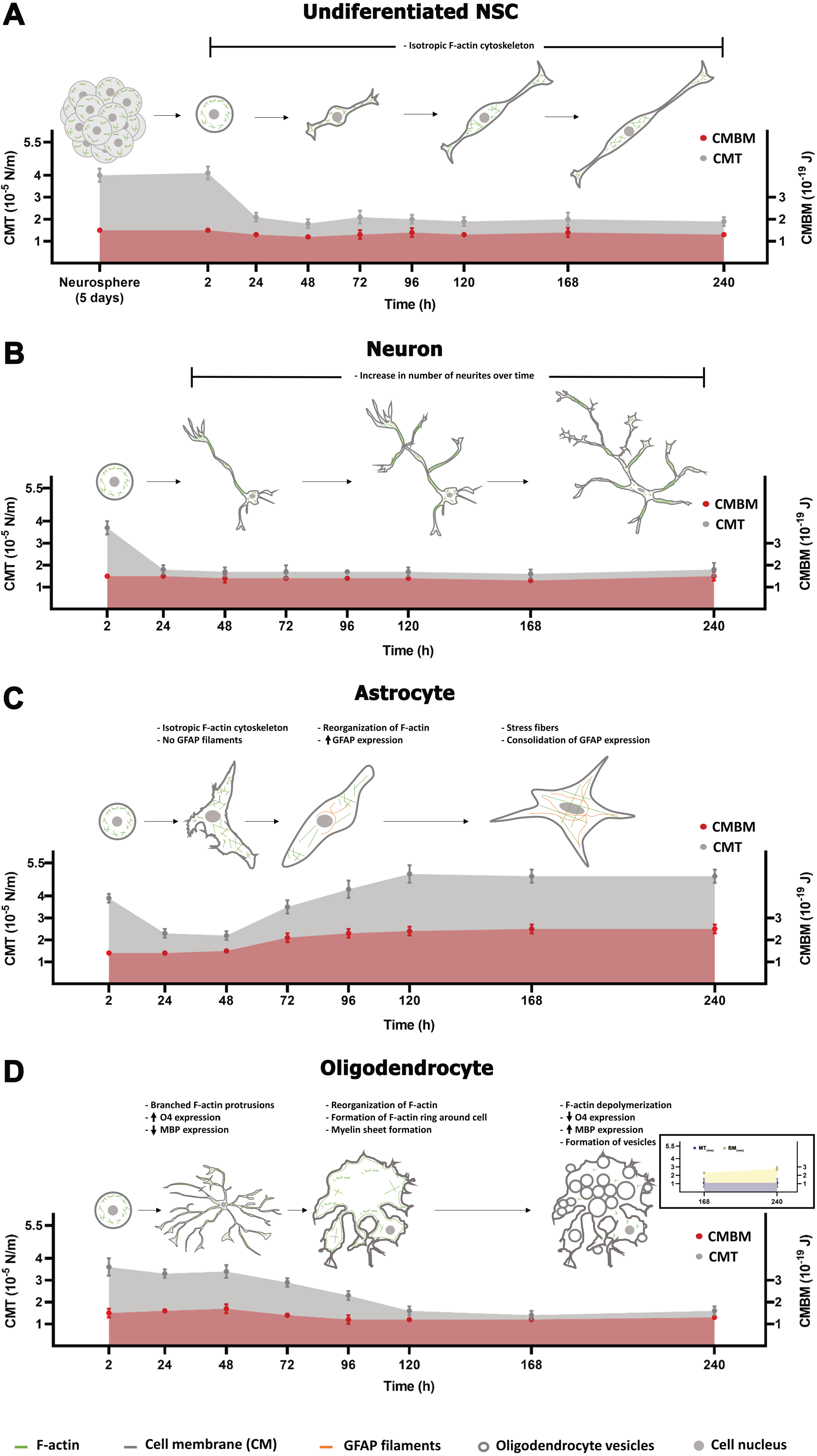
Schematic representation correlating the variations in morphological phenotype with CMT and CMBM over the course of (A) undifferentiated NSCs, (B) NSC differentiation into neurons, (C) NSC differentiation into astrocytes and (D) NSC differentiation into oligodendrocytes. Insert in D represents the membrane tension and bending modulus for oligodendrocyte vesicles.

In order to explore the elastic properties of these vesicles and to compare the results with those obtained from the oligodendrocyte surface, we next extracted tethers from these vesicles (Figure 6). Examples of the oligodendrocyte vesicles are shown in Figure 6A, with bright field images of three different situations: (image 3) when the bead is attached to the vesicle surface, (image 4) when the force is the maximum force *F*_*m*(*ves*)_ (before tether formation), and (image 5) when the tether is already formed and the measured force is the steady-state tether force *F*_0(*ves*)_. Figure 6B shows the force curve, with the numbered points corresponding to the numbered images in Figure 6A (images 3, 4 and 5). Figure 6C represents the mean values for *F*_*m*(*ves*)_ and *F*_0(*ves*)_ at 168 and 240h of differentiation. No differences among force values were observed for both time conditions. The values found were 14±2 pN and 15±2 pN for *F*_0(*ves*)_ and 50±12 pN and 53±13 pN for *F*_*m*(*ves*)_, respectively at 168 and 240h conditions. The bead/membrane contact patch radius, *R*_*p*(*ves*)_, was also measured by image analysis. Figure 6A (image 4) represents an example of a chosen frame and indicates how *R*_*p*(*ves*)_ was obtained in that case. *R*_*p*(*ves*)_ was measured for both conditions (168 and 240h). No statistically significant differences were observed (Figure 6D). The values found were 540±7 nm and 545±8 nm, respectively for the 168 and 240h conditions. Using the values of *R*_*p*(*ves*)_, *F*_*m*(*ves*)_ and *F*_0(*ves*)_, we calculated the values of the oligodendrocyte vesicles tether radius, *R*_(*ves*)_, using equation (2). The values found were 104±36 nm and 112±44 nm, respectively for the 168 and 240h conditions. Finally, the membrane tension, *MT*_(*ves*)_, and bending modulus, *BM*_(*ves*)_, for vesicles that appeared at the surface of oligodendrocytes at 168h and 240h were calculated.

Altogether, the results confirm that NSCs differentiate into oligodendrocytes; however, for this specific situation, the process seems to be slower than the others: although O4 appeared in the first hours of induction, MBP expression only appeared in very late hours. The change in actin cytoskeleton architecture, the increase in MBP expression and the formation of vesicles around the oligodendrocytes’ surface are all correlated with the decrease in CMT observed for these cells after 120h and until 240h (Figure 7D, grey curve and dots). The oligodendrocytes CMBM (Figure 7D, red curve and dots) values do not vary significantly. Finally, vesicles that appeared at the surface of oligodendrocytes at 168h and 240h present similar membrane tension values but higher bending modulus values when compared to those found for the oligodendrocyte cell surface (Figure 7D insert – yellow and blue curves and dots).

## DISCUSSION

The elastic properties of cell surfaces are increasingly recognized as key regulators of cell functions [1,14,16,43,44]. Here, we have mapped the variations in CM elastic properties during NSC differentiation into neurons, astrocytes and oligodendrocytes. The results show that these variations are correlated with morphological phenotype changes that can affect the differentiation process and ultimately the cell functions.

CMT has been considered an ideal candidate for global cell signaling, in view of its fast equalization around the cell in response to a stimulus [14], although controversies exist [45]. Conversely, CMBM is locally adjustable by a variety of mechanisms and is important for dynamic cell remodeling and movement [4]. These considerations emphasize the need to precisely determine CMT and CMBM for different cell types and in different biological contexts. Tether pulling experiments are a reliable choice method for such purpose [14]. However, it is important to measure not only the tether force but also the tether radius. Tether force can be measured with OT [6,7] or AFM [8,9]. However, measurement of tether radius is a challenge, as they are typically below the resolving limit of conventional optical microscopes. Some studies just assume a standard value for the CMBM (∼0.27 pN.µm – first obtained by [46]) and calculate CMT from direct measurements of tether force alone. This assumption that CMBM has a quasi-universal value for all cell membranes is unwarranted as CMBM itself is also dependent on tether force and radius [29,30]. Therefore, a correlative microscopy-based method previously established [7,13], was used in the present study. In this method, one extracts a tether and measures its force with OT and its radius with SEM. The intrinsic difficulty in performing the method is the main obstacle for this technique. Novel super-resolution live-microscopy techniques, capable of measuring simultaneously the tether radius and force, should be undertaken to move the field forward. Non-invasive methods have been recently developed [47,48] but still need further tests before they may be used as standards. Therefore, even if there might be some difficulties related with the correlative experiments performed in this work, they are still the gold-standard method to precisely determine the elastic properties of cell membranes.

Previous studies have performed tether extraction measurements from different stem cell types, such as human mesenchymal stem cells [49,50], mouse embryonic stem cells [51,52] and even mouse neural stem cells [53]. However, to the best of our knowledge, the present study is the first to measure both the steady-state tether force and tether radius for precise determination of the CM elastic properties of murine NSCs over the course of their differentiation into neurons, astrocytes and oligodendrocytes. We not only measured and mapped the changes in elastic properties of these cells during differentiation but also correlated these changes with morphological phenotype modifications acquired by these cells along this process.

Undifferentiated mouse NSCs used in this study have an embryonic origin and presented a low-density and isotropic actin network over the course of the entire experiment. This feature was previously documented for mouse embryonic stem cells [54] and conjectured to be necessary to maintain the balance between cortical and nuclear stiffness such that the rigidity of the cortex never exceeds the rigidity of the nucleus. This is important because, otherwise, the nucleus would end up sensing the cell’s own stiff cortex instead of the external mechanical environment. It is well established that nuclear stiffening follows stem cell differentiation [55]. The observed low-density isotropic actin network is correlated with low elastic constant values, as observed in the present study, suggesting that mouse NSCs have a weak interaction between the plasma membrane and the adjacent cortical cytoskeleton. Indeed, these cells must be prepared to give rise to the three cell types studied herein. Therefore, NSCs low CM elastic constants and less structured actin cytoskeleton can be important for allowing fast conversion to neurons, astrocytes and/or oligodendrocytes. More recently, researchers have shown in mice that NSCs can contribute to embryonic, early postnatal and adult neurogenesis in the hippocampus, and that these cells are continuously generated throughout a lifetime [56], which reinforces the importance of determining their CM elastic properties.

In our study we induced NSCs to differentiate into neurons. This induction occurred quickly and may be related to the period in which these cells were isolated, the neurogenesis window [17], with its peak happening between 12 and 16 days of embryonic development in mice. Thus, cells with neuronal phenotypes appeared in the first hours of differentiation and remained differentiated throughout the entire period, increasing the number and length of their protrusions. Interestingly, the CM elastic properties of neurons were also low and remained constant for almost the entire experiment. These mechanical features have been previousy described for differentiated mouse cortical and ganglionic eminence neurons [13]. The fact that these cells have low elastic constant values may be also related to a weak interaction between the plasma membrane and the cortical cytoskeleton.

Despite being practically constant in neurons and NSCs over the observerd period of this study, the CM elastic properties, particularly the CMT, changed dramatically in the first hours, decreasing to almost half its value before stabilization. This change in CMT is correlated with the cell spreading behavior that changes from initially round cells (grown in non-adherent spheres) to a final state in which the cells are fully spread/attached within the first 24 hours of experiment. Though never demonstrated before during NSCs differentiation, this behavior has been previously described for fibroblasts [57]. The CMT decreased during spreading and it was conjectured that the addition of new membrane and the increase in membrane area were mainly responsible for such phenomenon [57]. More recently, using embryonic stem cells, two different groups described, in quite recent studies yet to be published, that the decrease in CMT occurred when embryonic stem cells changed from their round and naïve state to a spread and primed state, and that the observed decrease in CMT was correlated with a decrease in membrane-cytoskeleton attachement [51,52] via GSK3β-driven β-catenin degradation [52], which in turn controls membrane tension and allows exit from naïve pluripotency. Our results are in agreement with all the above descriptions, even if the murine NSCs used in this study no longer display the transition between naïve and primed states. Moreover, all mentioned studies [51,52,57] assume a standard value for the CMBM and calculate CMT from direct measurements of tether force, without measuring the tether radius. As we have discussed, one should be careful when assuming that CMBM has a quasi-universal value for all cell membranes, since it depends on both tether force and radius [29,30]. In line with this, the results of the present study and previous observations [7,10,13] confirm that CMBM can vary depending on the cell type and cell context.

We also induced NSCs to differentiate into astrocytes. Although astrocytes share with neurons the same ectodermal origin, they are considerably more dynamic and mobile. Moreover, we noticed that the *in vitro* differentiation into astrocytes occurred more slowly, possibly owing to the fact that NSCs were isolated during the neurogenesis peak, between 12 and 16 days of murine embryonic development [17], while gliogenesis occurs only in late-gestation to perinatal periods [21]. Thus, cells with an astrocytic phenotype only appeared around 48-72 hours of differentiation after the consolidation of GFAP expression. The increase in CM elastic properties values followed the consolidation of GFAP expression and actin cytoskeleton reorganization that strikingly changed from a low-density isotropic meshwork to a more organized stress fiber pattern.

Lastly, we also induced NSCs to differentiate into oligodendrocytes. Similarly to what happened for astrocytes, the *in vitro* differentiation process also occurred more slowly for oligodendrocytes. This behavior partially recalls what happens during mouse embryonic development. Although oligodendrocyte progenitor cells are generated from NSCs in different locations and times during CNS development, the formation of mature myelinated oligodendrocytes only occurs around 5-7 days after birth in mice [38]. In our *in vitro* differentiation model, cells with the mature oligodendrocyte phenotype appeared around 120 hours after plating. This differentiation coincided with a decrease in O4 and an increase in MBP expression levels, and it correlated with a striking actin cytoskeleton reorganization, changing from initially actin-rich tubular protrusions to a more lamellar morphology, with actin moving towards the cell periphery until almost disappearing at later stages. This morphological behavior has already been documented in other studies [39-41]. Moreover, Nawaz et al. [40] performed tether pulling experiments in oligodendrocytes after being in culture for 48 hours (without myelin) and 120 hours (with myelin sheath). They found a ∼1.3-fold decrease in tether force, similarly to that found in our measurements. However, again, these authors did not measure the tether radius but concluded, only from the tether force, that the CMT was decreasing. This decrease was associated with actin depolymerization via ADF/cofilin and induced membrane spreading and myelin sheet growth [40]. We performed tether force and radius measurements for each experimental time point ranging from 2 to 240 hours of differentiation and demonstrated that indeed CMT was decreasing, while CMBM did not vary. Another feature that appeared in the last hours of differentiation (168 and 240 hours) was the presence of membrane vesicles at the oligodendrocytes surfaces. These vesicles were previously documented several years ago in ultrastructural caracterizations of cultured oligodendrocytes [42], but have been neglected since that time. The formation of these vesicles can be well explained based on the excess membrane production and the huge actin depolymerization associated with an increase in membrane-cortex detachment already described at latter stages of oligodendrocyte differentiation [39-41]. The elastic properties of these membrane vesicles were determined and compared with the elastic properties of the oligodendrocyte CM at these time points. The vesicle membrane tension was slightly lower when compared to the CMT but the vesicle bending modulus increased ∼2.3-fold when compared to the CMBM. These differences may be associated with the total decoupling of the plasma membrane from the underneath cortical cytoskeleton during vesicle formation. It is noteworthy that even with the increase in actin depolymerization, the plasma membrane of oligodendrocytes still has some connection with the cortical cytoskeleton, but this connection may be lost during vesicle formation.

In conclusion, we have measured the elastic constants of NSCs membranes over the course of their differentiation into neurons, astrocytes and oligodendrocytes. We have also presented experimental evidence supporting the conjectured correlation between cells’ surface elasticity and the functions of these cells during the differentiation process. All tether radii and forces were measured. The values, determined under uniform and controlled experimental conditions, not only reinforce confidence but can also be taken as data for future studies. The importance of CMT and CMBM is now established for a variety of cells. However, the mechanisms allowing cells to set and regulate their CM elastic properties remain to be elucidated. The present study contributes to the knowledge that the CM elastic constants not only differ among cell types, but also depend on the cell state at the time of measurement. A variety of cellular processes are already known to affect the CM elastic properties, but how they all manage to determine and maintain their values still remains a challenging question for the field.

## Acknowledgments

We acknowledge Dr. Barbara Hissa for critical reading and scientific editing of the manuscript. We also thank Dr. Grasiella Matioszek, Jefte Farias, Pedro Lourenço, Gabriela Maciel and the members of CENABIO electron microscopy facility for all-important help. This work was supported by the Brazilian agencies Conselho Nacional de Desenvolvimento Científico e Tecnológico (CNPq), Coordenação de Aperfeiçoamento de Pessoal de Nível Superior (CAPES) – Financial Code 001, and Fundação de Amparo à Pesquisa do Rio de Janeiro (FAPERJ). BP was supported by a JCNE grant from FAPERJ. NBV, HMN and BP are members of the Instituto Nacional de Ciência e Tecnologia de Fluidos Complexos. VMN is a member of the Instituto Nacional de Ciência e Tecnologia de Neurociência Translacional.

## Author Contributions

Conceptualization: LR, HMN and BP. Formal analysis: JS and BP. Funding acquisition: HMN and BP. Investigation: JS, GRSA, CF, DM and BP. Methodology: GRSA, MF, SF, NBV, LR and BP. Project administration: BP. Resources: VMN, MF, SF, NBV, LR, HMN and BP. Supervision: HMN and BP. Validation: JS, GRSA and BP. Visualization: BP. Writing – original draft: HMN and BP. Writing – review & editing: HMN and BP. All authors have read and agreed to the published version of the manuscript.

## Conflict of interest

The authors declare that they have no conflict of interest and no competing financial interest.

## Supplementary Materials

**<u>Figure S1:</u> Confirmation that NSCs remain undifferentiated in culture.** (A) Phase contrast image of neurospheres in culture and a representative neurosphere stained for nestin (in white) and sox-2 (in red), together with DAPI (in blue). Scale bars are respectively 100 µm and 50 µm. (B) Representative image of a dissociated neural stem cell (radial glia) after 240h in culture stained for nestin (in white) and sox-2 (in red) together with DAPI (in blue) and actin, with phaloidin-FITC (in green). Scale bar is 20 µm.

