## Supplementary figures and images for "Membrane Elastic Properties During Neural Progenitor/Neural Stem Cell Differentiation"

### Supplemental Figure 1

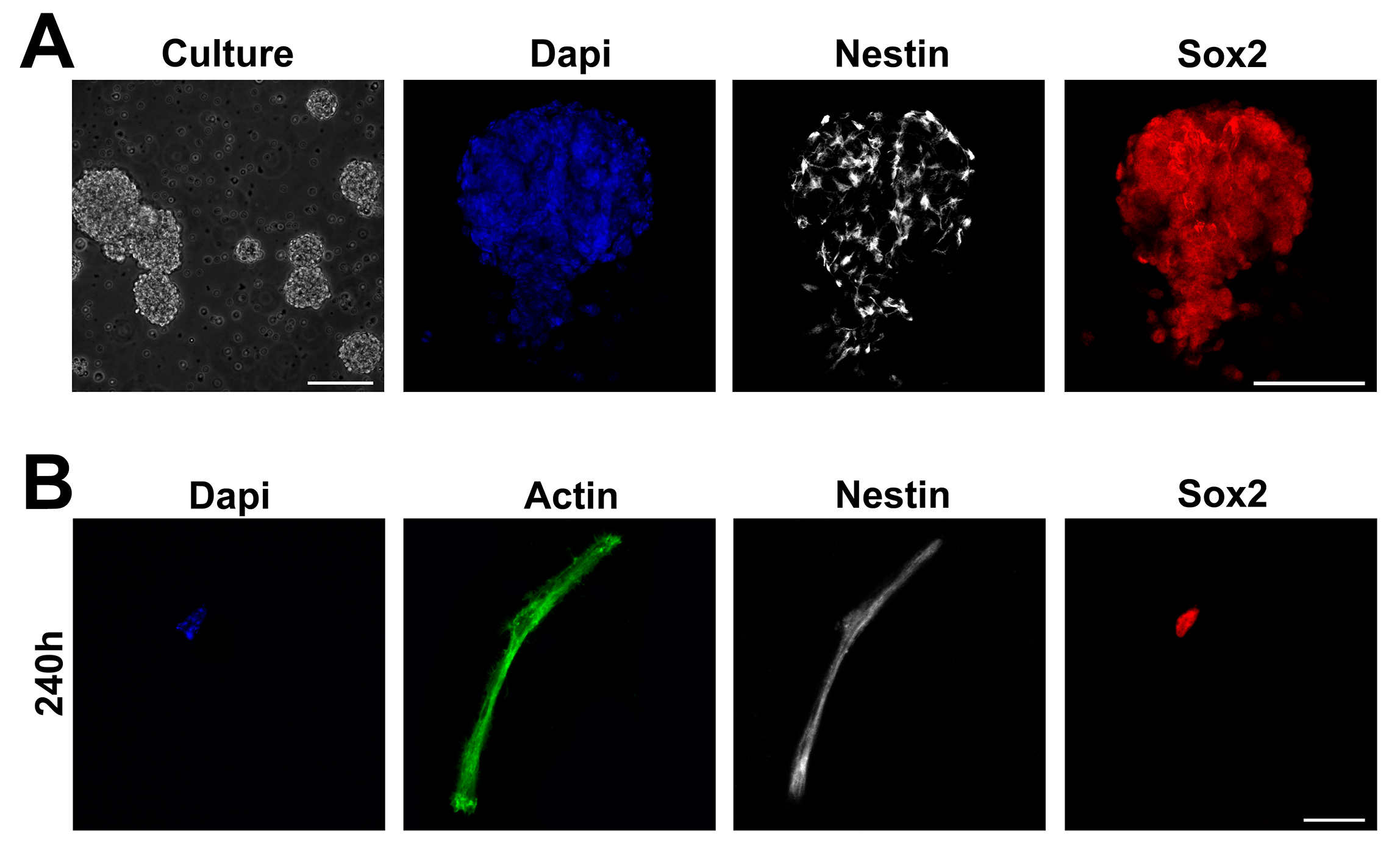
